# Coordinated infraslow cortical oscillations of neuromodulators during NREM sleep

**DOI:** 10.64898/2025.12.22.695982

**Authors:** Celia Kjaerby, Tessa Radovanovic, Eszter Rebeka Kovács, Zuzanna Bojarowska, Klaudia Anna Tokarska, Anastasia Tsopanidou, Elise Schiøler Nielsen, Mie Andersen, Yulong Li, Pia Weikop, Verena Untiet, Maiken Nedergaard

## Abstract

Neurotransmitters and neuromodulators regulate brain states through diverse mechanisms, yet how their activities are coordinated during sleep remains unresolved. Using *in vivo* fiber photometry in adult mice expressing genetically encoded fluorescent biosensors, combined with EEG/EMG recordings, we investigated the temporal organization of multiple neuromodulators during sleep in barrel cortex, with norepinephrine (NE) as a reference signal. All five neuromodulators examined, acetylcholine, serotonin, dopamine, histamine, and NE, exhibited synchronized infraslow cortical oscillations during NREM sleep. Optogenetic suppression of locus coeruleus (LC) neurons abolished NE oscillations and selectively reduced acetylcholine fluctuations in barrel cortex, whereas targeted inhibition of basal forebrain cholinergic neurons attenuated REM-associated acetylcholine elevations without disrupting NREM-related oscillations or NE dynamics. The synchronized infraslow cortical oscillations spanning multiple neuromodulators reveal a previously unrecognized mechanism for organizing sleep architecture.

## INTRODUCTION

Neuromodulators regulate brain states through distinct mechanisms, yet their coordinated activity during sleep remains poorly understood. Recent studies have demonstrated that the locus coeruleus (LC) exhibits periodic bouts of activity during NREM sleep, producing infraslow oscillations in extracellular norepinephrine levels approximately every 30-50 s. These oscillations play a critical role in organizing sleep microstructure, influencing spindle-dependent memory and glymphatic function^1–3^ and appear to be a global phenomenon based on reports from multiple regions^1,2,4,5^. Infraslow oscillations in pupil diameter (indirect measures of LC-norepinephrine) also occurs in humans during NREM sleep, where they inversely correlate with spindle clusters^6^, indicating a possible conserved neuromodulator rhythm across species.

Albeit, several other neuromodulators are reported to exhibit slow oscillatory patterns, inconsistencies in experimental design have made it difficult to determine whether these signals occur in synchrony. Given that infraslow fluctuations of norepinephrine induce slow vasomotion, resulting in large-amplitude cerebrospinal fluid (CSF) influx^3^, and that most other neuromodulators are vasoactive^3,7–10^, it is of considerable interest to investigate whether they exhibit coordinated oscillatory dynamics, potentially contributing to the global regulation of vascular tone and glymphatic function during sleep.

Many neuromodulators are involved in sleep-wake regulation^11,12^. For example, the pedunculopontine and laterodorsal tegmental nuclei (PPT and LDT) nuclei, as well as the basal forebrain, release acetylcholine, with the nucleus basalis of Meynert serving as the primary cortical source^13–17^. The raphe nucleus produces serotonin^18^, the ventral tegmental area releases dopamine^19^, and the hypothalamus is the source of histamine ^20,21^. While these neuromodulators generally play distinct but sometimes coactive roles in wakefulness^22,23^, emerging evidence suggests that they exhibit infraslow oscillations during NREM sleep. For instance, infraslow fluctuations of serotoninergic dorsal raphe neurons^24,25^ and extracellular serotonin^26^ and acetylcholine^27^ has been observed. However, differences in experimental designs - such as variations in recording techniques, brain regions studied, and temporal resolution - have hindered direct comparison^25,28–36^. Determining whether neuromodulators oscillate in synchrony has far-reaching implications - not only for understanding the microstructure of sleep - but also for uncovering the restorative functions of NREM sleep and for clarifying how their disruption may underlie sleep disturbances that precede, and potentially drive, the early pathogenesis of neurodegenerative diseases^8^. In this study, we demonstrate that acetylcholine, serotonin, dopamine, histamine, and norepinephrine - which all have vasoactive properties - exhibit coordinated infraslow oscillations in the barrel cortex during NREM sleep with a hierarchical organization in which LC–derived norepinephrine may serve a coordinating role in the generation of the neuromodulator oscillations. In conclusion, this study points to the possible existence of an integrated neuromodulatory network that orchestrates sleep microarchitecture and slow vasomotion, thereby promoting glymphatic clearance.

## RESULTS

### Neuromodulators display infraslow fluctuations during NREM sleep

To map the activity of norepinephrine, acetylcholine, serotonin, dopamine, and histamine during sleep, we performed bilaterally injections into the primary somatosensory barrel cortex (S1BF) using two viral constructs expressing fluorescent biosensors driven by the synapsin promoter, thereby targeting both excitatory and inhibitory neurons. Given the bi-hemispheric nature of sleep in mammals^37^, this approach allowed us to record the activities of two neuromodulators simultaneously during natural sleep. The norepinephrine biosensor, GRAB_NE2m_ ^29^, served as a reference, based on its well-established oscillating sleep patterns during sleep^1,2^. In the contralateral hemisphere, we injected AAV vectors carrying biosensors for either acetylcholine^34^, dopamine^31^, serotonin^28^ or histamine^32^ (Figure 1, Suppl. Figure S1A). Prior to data collection, we confirmed that the infraslow oscillatory pattern of norepinephrine in both hemispheres exhibited similar and tightly coupled dynamics (Suppl. Figure 1C). This setup allowed us to analyze the dynamic of each neuromodulator in relation to norepinephrine. To control for artifacts^38^, we simultaneously recorded 405 nm emission as a baseline for offline calculation of relative fluorescence changes (Suppl. Figure S1B). Norepinephrine-insensitive GRAB_NEmut_^1^ (and other neuromodulator controls^28,32,34^), green fluorescent protein (GFP) and simultaneous 405 nm control recordings displayed significantly smaller responses and narrower distributions compared to the norepinephrine signal, confirming the signal-specific fluctuations of the GRAB_NE2m_ biosensor (Suppl. Figure S1D-F).

**Figure 1.**
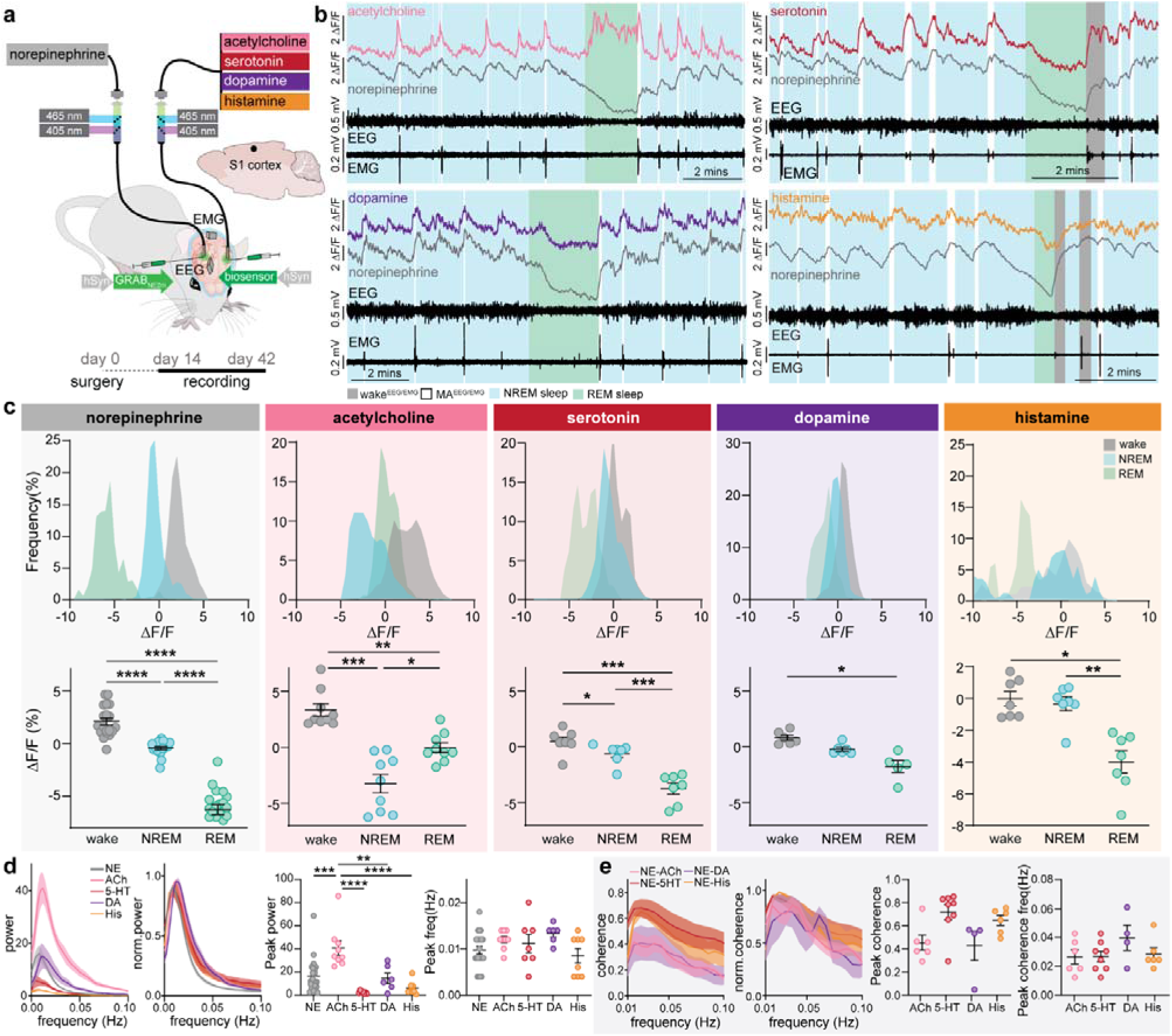
All main neuromodulators display infraslow fluctuations during NREM sleep. (A) The norepinephrine sensor, GRAB_NE2m_, was expressed in one hemisphere of the primary somatosensory barrel cortex (S1BF) and a biosensor for either acetylcholine, dopamine, serotonin or histamine were expressed on the contralateral side alongside EEG and EMG electrodes. Animals were implanted with optic fibers above both hemispheres of barrel cortex to image norepinephrine (grey) simultaneous with either acetylcholine (pink), serotonin (red), dopamine (purple) or histamine (orange). (B) Representative traces of simultaneous recordings with EEG/EMG-based sleep scoring in acetylcholine, serotonin, dopamine and histamine expressing animals. MA: micro-arousal. (C) Frequency distribution of ΔF/F values during wake, NREM sleep and REM sleep for each biosensor as well as summary plots (RM one-way ANOVA or Mixed-effects model with Tukey’s multiple comparisons test). (D) Power of infraslow oscillations of neuromodulators during NREM sleep as well as normalized power and summary plots of peak power and peak frequency (one-way ANOVA with Tukey’s multiple comparisons test). (E) Coherence of norepinephrine and the other neuromodulator. Also shown is normalized coherence as well as peak power and frequency (one-way ANOVA with Tukey’s multiple comparisons test). Data is shown as mean±SEM. Norepinephrine: n = 20 (histogram), n = 22 (power); acetylcholine: n = 9 (histogram+power), n = 6 (coherence); serotonin: n = 7 (histogram+power), n = 8 (coherence); dopamine: n = 6 (histogram+power), n = 4 (coherence); histamine: n = 7 (histogram), n = 8 (power), n = 6 (coherence). \**P*< 0.05, \*\**P*< 0.01.\*\*\**P*< 0.001, \*\*\*\**P*< 0.0001.

By conducting continuous recordings spanning multiple sleep-wake episodes, we observed a prominent infraslow oscillatory pattern not only for norepinephrine but also for other neuromodulators during NREM sleep (Figure 1). The distribution of ΔF/F values for each biosensor during wake, NREM and REM sleep revealed distinct dynamics: norepinephrine, acetylcholine and serotonin were reduced during NREM sleep compared to wakefulness, whereas dopamine and histamine showed no difference (Figure 1C). Notably, the lowest neuromodulator levels were observed during REM sleep with acetylcholine being the only exception. As expected, acetylcholine levels were higher during REM than NREM sleep but still lower than during wakefulness^39^.

To further investigate the infraslow oscillations, we performed a power analysis (Figure 1D). All neuromodulators exhibited infraslow oscillations within the same frequency range, peaking at ∼ 0.01 Hz. It is important to note, however, that GRAB biosensors vary considerably in fluorescence intensity, binding kinetics, and ligand affinity, which makes direct comparison of *in vivo* signals across neuromodulators challenging^25,28–36^. To mitigate these limitations, we restricted our cross-modality comparisons to dynamic changes relative to norepinephrine.

To assess synchronization, we conducted a coherence analysis between the reference norepinephrine signal and the other neuromodulators. Coherence peaked at ∼0.02 Hz (Figure 1E) with no difference in peak frequency (Figure 1E) between the neuromodulators highlighting the synchronized nature of these oscillations in the infraslow range.

### Neuromodulators share coordinated infra-oscillatory fluctuations during sleep state transitions

Next, each norepinephrine ascent during NREM sleep was classified into three distinct types: 1) ascent not associated with an EMG fluctuations (Figure 2A, termed NREM-NREM, detailed discussion ^40^), 2) micro-arousals with evident EMG events (lasting < 15 seconds, referred to as MA^EEG/EMG^) and 3) wakefulness (EMG events lasting > 15 seconds, referred to as wake^EEG/EMG^). These states were characterized by increasing arousability, as reflected by desynchronization in the delta, theta, sigma and beta frequency bands, along with elevated low- and high-gamma power, progressing from NREM-NREM to MA^EEG/EMG^ to wake^EEG/EMG^ (Suppl. Figure S2).

**Figure 2.**
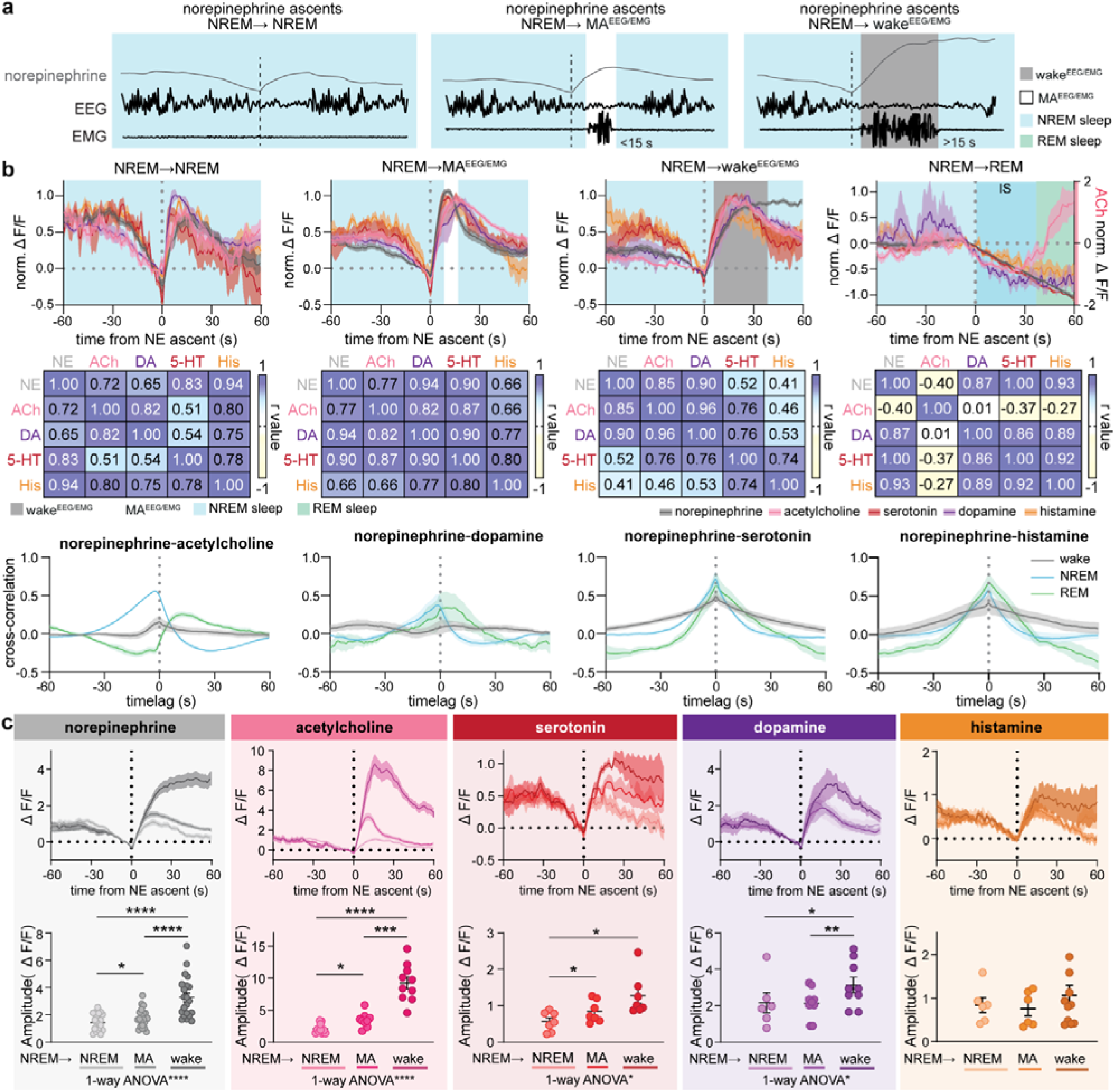
Neuromodulators share coordinated infra-oscillatory fluctuations during sleep state transitions. (A) Illustrative traces showing how norepinephrine (NE) ascents were categorized into three behavioral outcomes: No EMG events (termed NREM-NREM transition), EMG events lasting <15 s (termed NREM-micro-arousal (MA^EEG/EMG^)) and EMG events >15 s (termed NREM-wake^EEG/EMG^). (B) Mean traces normalized to span from 0 to 1 across NREM to NREM, NREM to MA^EEG/EMG^, NREM to wake^EEG/EMG^ and NREM to REM sleep with corresponding correlation plots below (Pearson r values shown. NE, norepinephrine; ACh, acetylcholine; DA, dopamine; 5-HT, serotonin; His, histamine. All correlations have p-values below 0.0001 except acetylcholine-dopamine during NREM-REM transitions (P = 0.799)). Cross-correlations between norepinephrine and each neuromodulator are shown during NREM, REM and wake. (C) ΔF/F mean traces for each biosensor aligned to onset of NE ascent (and zeroed) at transitions from NREM to wake (dark color), NREM to MA^EEG/EMG^ (medium color) and NREM to NREM (light color). Beneath is shown the amplitude (change from before (-10 to 0 s) to after (10-30 s)) for each biosensor for each transition. 1-way ANOVA was performed to test differences across arousal transitions. Norepinephrine, n = 21; acetylcholine, n = 10; serotonin, n = 7; dopamine, n = 6; histamine, n = 4. \**P*< 0.05, \*\**P*< 0.01, \*\*\*\**P*< 0.0001.

To compare all neuromodulators dynamics across arousal transitions, we normalized the amplitude of neuromodulator rises (aligned to norepinephrine ascent) to account for potential differential biosensor sensitivities (Figure 2B). The analysis revealed similar fluctuations in acetylcholine, serotonin, dopamine, and histamine across states (summary plots shown in Suppl. Figure S3). High correlations (Figure 2B, middle) and positive cross-correlations during NREM sleep (Figure 2B bottom) confirmed their synchronization. During wakefulness, cross-correlations remained positive but weaker. During REM sleep, strong positive correlations were evident for all neuromodulators except acetylcholine, which exhibited an inverse correlation due to its increase during this state. Previous studies have shown that norepinephrine levels begin to descend prior to REM sleep onset during a type of spindle-rich NREM phase known as intermediate state sleep; REM sleep onset typically occurs around 40-60 seconds after norepinephrine starts to descend^1,41^. We validated the spectral properties of this period (Suppl. Figure S2 right) and found that acetylcholine, serotonin, dopamine, and histamine also exhibited descending trends during intermediate state sleep (Figure 2B), which continued into REM sleep. However, acetylcholine uniquely began to ascend monotonically upon REM sleep onset consistent with prior findings^42,43^ (Figure 2B). Lastly, we analyzed the differential neuromodulator increases during arousal transitions. As previously described^1^, the amplitude of the norepinephrine ascents were significantly associated with behavioral outcomes: ascents associated with awakenings were largest, followed by those linked to micro-arousals and then norepinephrine ascents without EMG events. Similar patterns were also observed for acetylcholine, serotonin and dopamine, with histamine as the exception (Figure 2C). While histamine displayed small ascents during transitions, these were less pronounced and did not vary with arousal levels (Figure 2C). Only during extended awakenings did histamine levels begin to rise prominently (Suppl. Figure S4). Such temporal dynamics likely underlie the previously observed lack of difference in mean histamine levels between wake and NREM sleep (Figure 1C), given that recordings were conducted during the light phase, when wake episodes are relatively short.

### Optogenetic suppression of the locus coeruleus reduces micro-arousals

The preceding results suggest a coordinated temporal rhythm across multiple neuromodulatory systems during NREM sleep. However, the hierarchical relationship among these systems remains unclear. Given the well-established role of the LC–norepinephrine system as a key regulator of arousal and sleep–wake dynamics, we next examined how manipulating LC activity influences other neuromodulatory systems and behavioral outcome, exemplified by cortical acetylcholine release and the occurrence of micro-arousals. To explore the impact of norepinephrine suppression on NREM and REM sleep-dependent acetylcholine dynamics, we first employed optogenetic inhibition of LC. We introduced either the light-responsive proton pump, eArch3.0-eYFP (Arch), or an eYFP-only control construct into TH-positive neurons of both LC hemispheres. GRAB_NE2m_ and GRAB_ACh3.0_ were expressed in opposite cortical hemispheres, to monitor norepinephrine and acetylcholine, respectively. Following these injections, we implanted fiber optic cannulas as well as screws for EEG and EMG measurements, as described previously^1^ (Figure 3A). We employed a real-time norepinephrine decline (threshold value of -5 ΔF/F (%)) to trigger a 2 min optogenetic LC suppression (Figure 3A), which resulted in LC suppression predominantly occurring during NREM sleep (Suppl. Figure S5C). The number of MA^EEG/EMG^ events was reduced when LC activity was inhibited, consistent with prior observations (Figure 3B)^1^. Importantly, the optogenetic suppression protocol did not cause significant changes in overall sleep structure (Suppl. Figure S5B-C). We next posed the reverse question: does LC stimulation induce microarousals? Optogenetic activation of LC (2 s, 10 ms, 20 Hz 465 nm pulses) during NE descends produced MA^EEG/EMG^ in ∼80% of the stimulations compared to 15% in YFP-expressing control animals during NREM sleep (Suppl. Figure S6A-C). Thus, stimulation of LC is capable of inducing microarousals. Combined, the optogenetic manipulation of LC suggests that norepinephrine plays an important role in controlling the frequency of microarousals.

**Figure 3.**
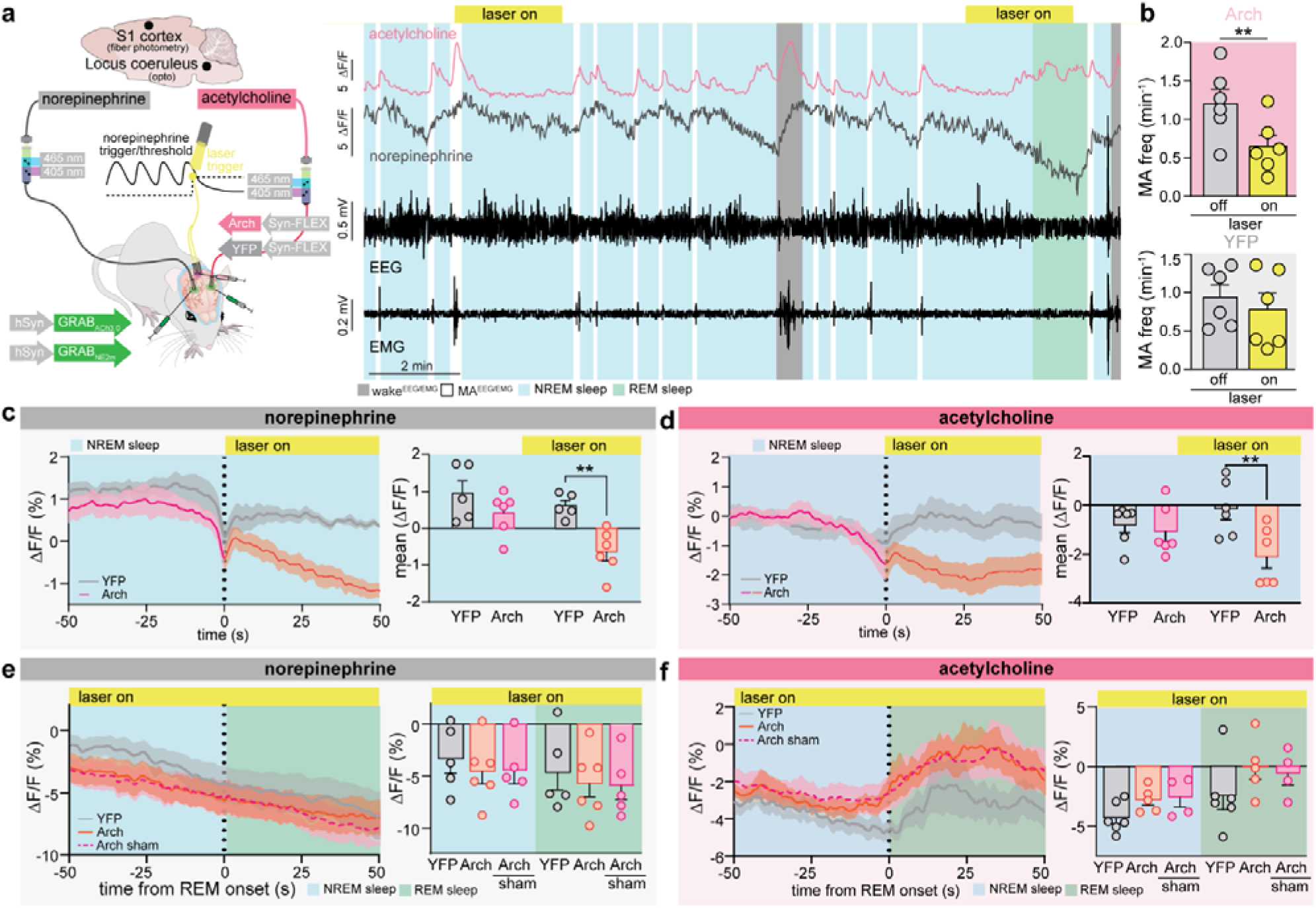
Optogenetic suppression of the locus coeruleus reduces micro-arousals. (A) Arch was expressed in locus coeruleus (LC) of TH::Cre mice, while GRAB_NE2m_ and GRAB_ACh_ were expressed in barrel cortex. Green laser light triggered by norepinephrine decline (closed loop) was delivered to LC during combined fiber photometry and EEG/EMG recordings. Example traces showing the effects of 2 min optogenetic suppression of LC on norepinephrine and acetylcholine levels are displayed. (B) Frequency of EMG-defined micro-arousals (MA) with and without optogenetic LC suppression. n = 6 Arch, 6 YFP. MA: micro-arousal (Paired t test). (C) Mean norepinephrine (n = 6 Arch, 5 YFP) and (D) acetylcholine (n = 6 Arch, 6 YFP) traces during NREM sleep aligned to laser stimulation onset in Arch and YFP-expressing animals. Summary plot showing before (-10 to 0 s) and after (20 to 30 s) laser onset. (E) Mean norepinephrine and (F) acetylcholine traces during NREM-REM sleep transition aligned to REM sleep onset within the first 60 s of laser on (Arch, NE/ACh: n = 6/5; YFP, NE/ACh: n = 5/6) and REM sleep episodes during laser off (Arch, NE/ACh: n = 5/4). Summary plot showing before (-10 to 0 s) and after (20 to 30 s) REM sleep onset (2-way repeated measures ANOVA (sleep stage: NE***, ACh**)). n = 6 Arch, n = 6 YFP (NE, n = 5 due to poor expression). Data is shown as mean±SEM. \**P*< 0.05, \*\**P*< 0.01, \*\*\**P*< 0.001.

Acetylcholine exhibits ascents during both NREM (associated with micro-arousals and awakenings^15,44–46^) and REM sleep. We next asked whether LC suppression is capable of inhibiting acetylcholine oscillations during NREM sleep. We focused the first analysis on norepinephrine and acetylcholine levels within the first 50 s after laser onset to target NREM over REM sleep (Figure 3C-D). We found that LC suppression induced a consistent reduction in both norepinephrine and acetylcholine levels compared to YFP controls (Figure 3C-D, for spectral properties see Suppl. Figure S7A). Thus, optogenetic inhibition of the LC suppresses acetylcholine oscillations during NREM sleep, possibly as a downstream consequence of the reduction in LC-driven micro-arousals.

Can LC suppression during REM sleep also suppress the rise in acetylcholine? To address this question, we identified REM sleep episodes with the laser on (Arch) versus off (Arch-sham) within the same animal and compared to the YFP-control group. As expected, norepinephrine levels decline during REM sleep, consistent with LC quiescence^1,41^ (Figure 3E-F). LC suppression did not further reduce norepinephrine levels and acetylcholine increases during REM sleep regardless of laser or Arch/YFP status (Figure 3E-F, for spectral properties see Suppl. Figure S7B-C). These findings suggest that LC-norepinephrine activity modulates acetylcholine release during NREM, but not during REM sleep.

### Optogenetic suppression of the cholinergic basal forebrain does not impact micro-arousals

Next, we asked whether suppression of the cholinergic basal forebrain could similarly inhibit neuromodulator-driven oscillations, as observed with LC suppression. To test this, we injected Cre-dependent eArch3.0 into the nucleus basalis of Meynert in Chat-Cre mice and recorded norepinephrine and acetylcholine signals in primary somatosensory barrel cortex (Figure 4a, Suppl. Figure S5D-F). We confirmed that basal forebrain suppression during REM sleep, when acetylcholine levels typically rise, reduced acetylcholine levels compared to laser-off (Arch-sham) control conditions (Figure 4C, for spectral properties see Figure S7E-F). This effect was most pronounced at the end of the stimulation. In contrast, norepinephrine declined steadily throughout REM sleep and was unaffected by basal forebrain suppression (Figure 4D). During NREM sleep, basal forebrain suppression did not alter acetylcholine or norepinephrine levels, nor did it alter MA^EEG/EMG^ frequency or overall sleep composition (Figure 4B, E-F, Suppl. Figure S5E).

**Figure 4.**
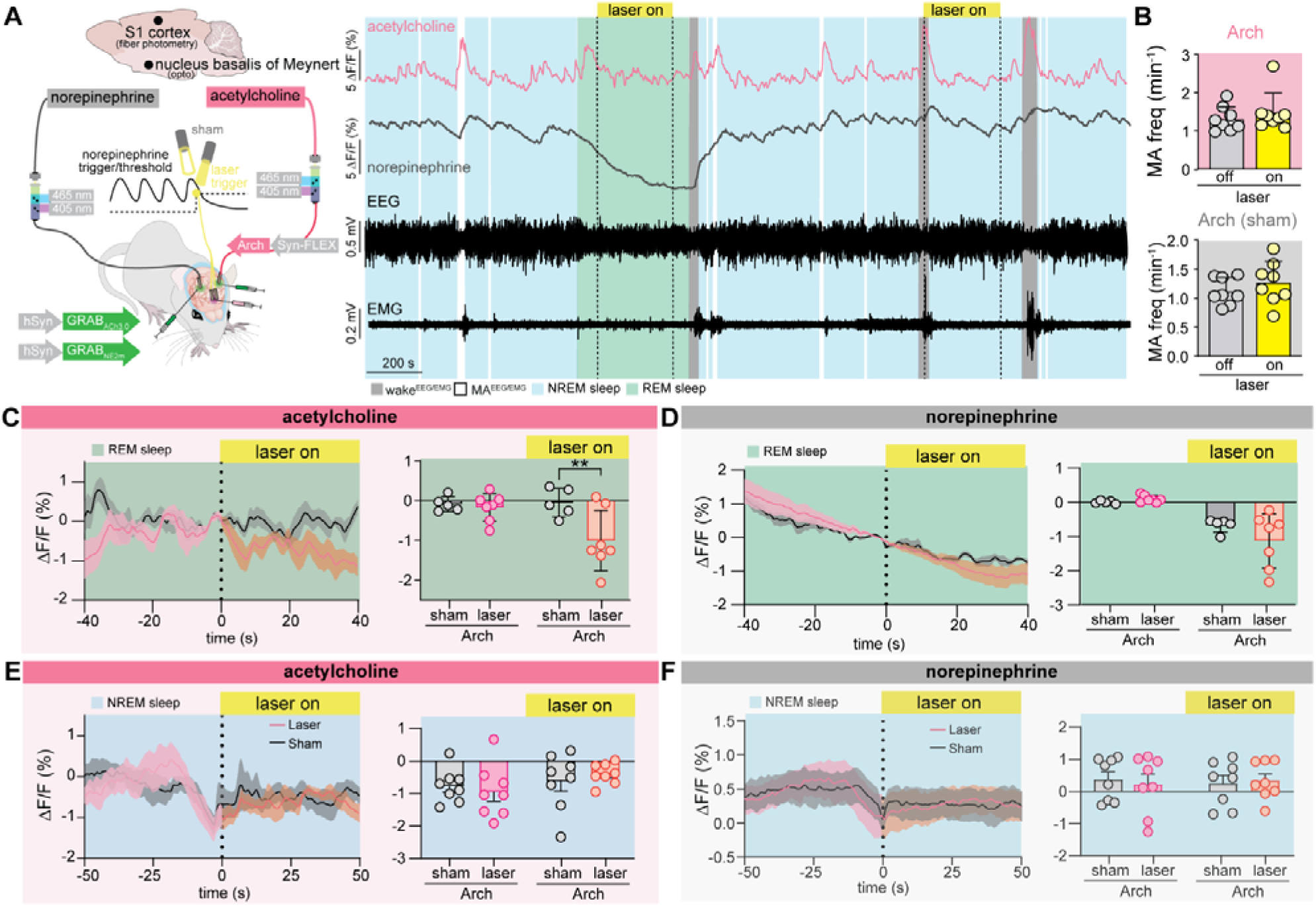
Optogenetic suppression of the cholinergic basal forebrain does not impact micro-arousals. (A) Arch was expressed in basal nucleus of Meynert (NBM) in Chat:Cre mice, while GRAB_NE2m_ and GRAB_ACh_ were expressed in barrel cortex. Green laser light triggered by norepinephrine decline (closed loop) was delivered to basal forebrain. Example traces showing the effect of optogenetic suppression of basal nucleus of Meynert on norepinephrine and acetylcholine levels are shown. (B) Frequency of EMG-defined micro-arousals (MA) with and without suppression of basal forebrain. n = 8 Arch. MA: micro-arousal (Paired t test). (C) Mean acetylcholine and (D) norepinephrine traces showing effect of laser onset on neuromodulator levels during REM sleep only (before-after × laser on/off*, Arch, ACh/NE: n = 7/5). Summary plot showing before (-10 to 0 s) and after (30 to 40 s) laser onset (2-way repeated measures ANOVA (time x condition: ACh*, NE)). (E) Mean acetylcholine and (F) norepinephrine (n = 8 Arch) traces during NREM sleep aligned to NE threshold crossing with or without laser stimulation of basal forebrain of Meynert in Arch-expressing animals. Summary plot showing before (-10 to 0 s) and after (30 to 40 s) laser onset (2-way repeated measures ANOVA (time x condition: ACh, NE)). Data is shown as mean±SEM. \**P*< 0.05, \*\**P*< 0.01, \*\*\**P*< 0.001.

## DISCUSSION

In this study, we demonstrate that infraslow cortical oscillations during NREM sleep are not exclusive to norepinephrine. Rather, norepinephrine, acetylcholine, serotonin, dopamine, and histamine all exhibit synchronized oscillatory dynamics in the cortex. While previous studies have independently documented infraslow fluctuations in acetylcholine, serotonin and dopamine levels in various brain regions^25,27,30–32,34,35,47–50^, differences in experimental design, such as targeting distinct brain areas or using divergent recording methodologies, have thus far precluded a direct, multiple-neuromodulator comparison. Notably, we also found that the amplitude of these neuromodulator oscillations scales with arousal level. Thus, the infraslow synchronized activity of cortical norepinephrine, acetylcholine, serotonin, dopamine, and histamine suggests a coordinated neuromodulatory rhythm during sleep.

The orderly synchronized oscillations of neuromodulators during NREM sleep starkly contrast with their heterogeneous and often specialized roles during wakefulness^51,52^. In the awake brain, norepinephrine, dopamine, serotonin, acetylcholine, and histamine regulate a wide array of distinct cognitive and behavioral functions, including emotion, reward and motivation, attention, sensory processing, and motor coordination^53,54^. During wakefulness, the neuromodulatory systems often operate in a dissociated manner^22^, but dependent upon the task can also be engaged in a coordinated manner. For example, orexin and norepinephrine can play coactive roles in wakefulness^23^. Yet, the emergence of highly synchronized infraslow oscillations across these systems during sleep may suggest a global, state-dependent reorganization of neuromodulatory tone to accommodate internal regulation and repair^55–57^. One possibility is that these unified oscillatory dynamics represent a coordinated recruitment of neuromodulators for homeostatic, autonomic, or restorative functions unique to sleep. Supporting this view, infraslow norepinephrine oscillations have been shown to temporally structure sleep microarchitecture by modulating sleep spindle density and glymphatic function^1,3,6,58,59^.

Next, we investigated the relative contributions of norepinephrine and acetylcholine to each other’s oscillatory dynamics. Our observation showed that optogenetic activation of LC increased micro-arousal frequency, whereas LC inhibition suppressed oscillation of both norepinephrine and acetylcholine and reduced the frequency of micro-arousals. LC is anatomically and functionally extensively linked to the other neuromodulatory systems in the brain. It sends direct projections to the serotonergic dorsal raphe, the pontine cholinergic PPT/LDT nuclei, the basal forebrain cholinergic nucleus basalis of Meynert, the dopaminergic ventral tegmental area, and histamine-releasing hypothalamus allowing LC to enable the activity of those other neuromodulator neurons^60^. Interestingly, LC suppression did not affect acetylcholine levels or EEG spectral features during REM sleep, indicating that cholinergic regulation during REM sleep relies on distinct, LC-independent pathways. Manipulation of the nucleus basalis of Meynert were able to lower the REM sleep specific increase of acetylcholine but did not alter norepinephrine during NREM sleep. Moreover, because LC suppression reduces MA^EEG/EMG^ events and LC activation increases micro-arousal frequency, the LC appears more likely than acetylcholine to initiate NREM sleep-imbedded micro-arousals^1–3,61^. While these observations point to that LC, rather than the nucleus basalis of Meynert, plays a dominant role in regulating neuromodulator oscillation during NREM sleep, additional studies are needed.

One point of attention is that all of the neuromodulators studied here are potent vasoactive compounds, triggering either constriction (norepinephrine, serotonin) or dilation (acetylcholine, dopamine) of the cerebral vasculature^7,10^. Histamine, which has mixed vascular effects^9^, displayed the smallest oscillation amplitude among the neuromodulators. We have recently shown that norepinephrine-induced vasomotion drives glymphatic flow during NREM sleep^3^. The significance of these findings lies in the emerging understanding that glymphatic clearance failure is a critical contributor to the accumulation of neurotoxic proteins such as amyloid-β, tau, and α-synuclein, a hallmark of neurodegenerative diseases, supported by both preclinical and clinical studies^62,63^. In a mouse model of Alzheimer’s disease, for instance, infraslow fluctuations of norepinephrine were disrupted following sleep deprivation, leading to impaired clearance and accumulation of insoluble amyloid-β^8^. Given that acetylcholine, serotonin, dopamine, and histamine are vasoactive, we suggest that the synchronized fluctuations of neuromodulators may collectively contribute to this vascular-coupled mechanism of sleep-dependent brain maintenance. These neuromodulatory systems in barrel cortex may work in concert to fulfill a shared physiological function: orchestrating brain-wide rhythms that promote synaptic homeostasis, memory consolidation, and the removal of metabolic waste during sleep.

### Limitations of the study

The recordings in this study were confined to the barrel cortex, and therefore the coordinated neuromodulator activity we report reflects dynamics within this cortical region. Numerous studies using diverse methodologies have demonstrated that norepinephrine, acetylcholine, serotonin, histamine, and dopamine are all released within barrel cortex^64–67^. Our findings show that each of these systems exhibits infraslow oscillations here. Comparable rhythms have been observed in other cortical and subcortical areas, raising the possibility that this phenomenon represents a broader, brain-wide rhythm. However, simultaneous measurements across multiple regions will be needed to conclusively establish its global nature.

Our study was not designed to test the causal contribution of each neuromodulatory system to those slow dynamics. Although our analyses suggest that the LC may play a dominant and potentially hierarchical role, additional systems are likely involved in initiating and sustaining the rhythm. Orexin (hypocretin) is a compelling candidate, given its key function in arousal control and its direct, excitatory projections to multiple neuromodulatory centers, including the LC^49,68^. The limited specificity of currently available genetic tools for manipulating orexin neurons remain a constraint^69–71^ preventing definitive test of its role in coordinating these rhythms.

## Supporting information

Supplementary Tabe S1

## RESOURCE AVAILABILITY

### Lead contact

Further information and requests for resources should be directed to and will be fulfilled by the lead contact, Maiken Nedergaard.

### Materials availability

The study did not generate new unique reagents.

### Data and code availability

- The datasets generated during and/or analyzed in this study are available from the corresponding author upon reasonable request.
- The custom-made MATLAB code used in this study is available from GitHub: https://github.com/MieAndersen/NE-oscillations
- Any additional information required to reanalyze the data reported in this paper are available from the lead contact upon request.

## ACKNOWLEDGMENT

We would like to thank Dan Xue for expert graphical support, This work was supported by Lundbeck Foundation grants R386-2021-165 (M.N.) and R413-2022-622 (C.K.); Novo Nordisk Foundation grants NNF20OC0066419 (M.N.); National Institutes of Health grants R01A 012707 and AT011439 (M.N.); National Institutes of Health grant U19NS128613 (M.N.); US Army Research Office grant MURI W911NF1910280 (M.N.); Human Frontier Science Program grant RGP0036 (M.N.); the Dr. Miriam and Sheldon G. Adelson Medical Research Foundation (M.N.); Simons Foundation grant 811237 (M.N.); and Cure Alzheimer Fund, Danmarks Frie Forskningsfond grants 3101-00282B (M.N.) and 2100-00018B (C.K.), JPND/HBCI 1098-00030B (M.N.), and JPND/Good Vibes 2092-00006B (M.N.).

## AUTHOR CONTRIBUTIONS

Conceptualization: C.K., V.U. and M.N. Fiber photometry and EEG methodology and investigation: C.K., T.R., E.R.K., K.T., Z.B., E.S.N, M.A., P.W. and A.T. Viral constructs: Y.L. Analysis: C.K., T.R., E.R.K., K.T. Z.B. and A.T. Visualization: C.K., T.R., E.R.K. and K.T. Supervision: M.N. and C.K. Writing—original draft: C.K., T.R., E.R.K. and M.N. Writing—review and editing: C.K., T.R., E.R.K. and M.N.

## DECLARATION OF INTERESTS

M.N. is a paid consultant for CNS2 for unrelated work.

## STAR Methods

### EXPERIMENTAL MODEL DETAILS

Wild-type C57BL/6 mice, aged 7 weeks, were obtained from Janvier Labs. Heterozygous TH::Cre (B6.Cg-7630403G23RikTg(Th-cre)1Tmd/J, Jackson Laboratory) and Chat:Cre (B6;129S6-Chattm2(cre)Lowl/J, Jackson Laboratory) mice were bred on a C57BL/6 background. Male and female mice of both genotypes were included in the study. The mice were housed under normal conditions with unrestricted access to food and water. The housing environment maintained a 12-hour light/dark cycle at a temperature of 21°C with a humidity range of 40-60%. The mice used for sleep assessments were between 12 and 20 weeks old. All experimental procedures were conducted in accordance with the guidelines set forth by the Danish Animal Experiments Inspectorate and were overseen by the University of Copenhagen Institutional Animal Care and Use Committee, adhering to the legislation outlined in the European Communities Council Directive of 22 September 2010 (2010/63/EU) concerning the welfare of animals used for scientific purposes.

## METHOD DETAILS

### Surgery

All surgical procedures were conducted in compliance with institutional guidelines. The mice used for surgery were aged between 7 and 15 weeks. General anesthesia was administered using 5% isoflurane, with maintenance levels set at 1-3% isoflurane. After placing the mice in the stereotactic frame, pre-operative buprenorphine was administered (0.05 mg kg^−1^) for general analgesia, along with lidocaine (0.03 mg kg^−1^) at the incision site. A midline incision was made on the scalp between the ears, and the skull was properly aligned. Using stereotactic coordinates relative to bregma, six burr holes were drilled in the skull using an electrical drill (Tech 2000, RAM Microtorque). In order to achieve strong infraslow oscillations^59^, we chose to target the primary somatosensory barrel cortex (S1BF), where we injected AAV9-hSyn-GRAB_NE2m_. In the contralateral barrel cortex, animals received one of the following biosensors: 1) hSyn-GRAB_NE2m_^29^ (norepinephrine, control, EC_50_/norepinephrine 380 nM, EC_50_/dopamine 20 µM), 2) hSyn-GRAB_ACh3.0_^34^ (acetylcholine), 3) hSyn-GRAB_5HT2h_^28^ (serotonin), 4) hSyn-GRAB_DA3h_^31^ (dopamine; EC_50_/dopamine 12 nM, EC_50_/norepinephrine 660 nM), or 5) hSyn-GRAB_HA1h_^32^ (histamine; Table 1). The stereotactic coordinates for barrel cortex injections were A/P -1.7 mm, M/L +/− 3.0 mm, and D/V −1.20 mm, −1.35 mm, −1.50 mm (200 nl of virus infused at each depth). Virus infusion was carried out at a rate of 100 nl min^−1^, and the needle remained in place for an additional 7 minutes before being slowly withdrawn. Following this, low-impedance stainless steel screws (0.8 mm, NeuroTek) were carefully secured into two burr holes located above the frontal cortex and above the cerebellum (serving as a reference area). Two silver wires (W3 Wire International) were inserted into the trapezius muscle to serve as EMG electrodes. Mono fiberoptic cannulas (400 μm, 0.48 NA, Doric Lenses or RWD Life Science), attached to a 2.5 or 1.25 mm-diameter metal ferrule, were then implanted bilaterally in the barrel cortices (A/P -1.7 mm, M/L +/− 3.0 mm, D/V −1.30 mm). Cannulas and screws were firmly affixed to the skull using dental cement (SuperBond). The animals received carprofen (5 mg kg^−1^) prior to regaining consciousness. A minimum recovery period of 2 weeks was allowed to ensure sufficient expression and overall well-being of the mice.

**Table 1.**
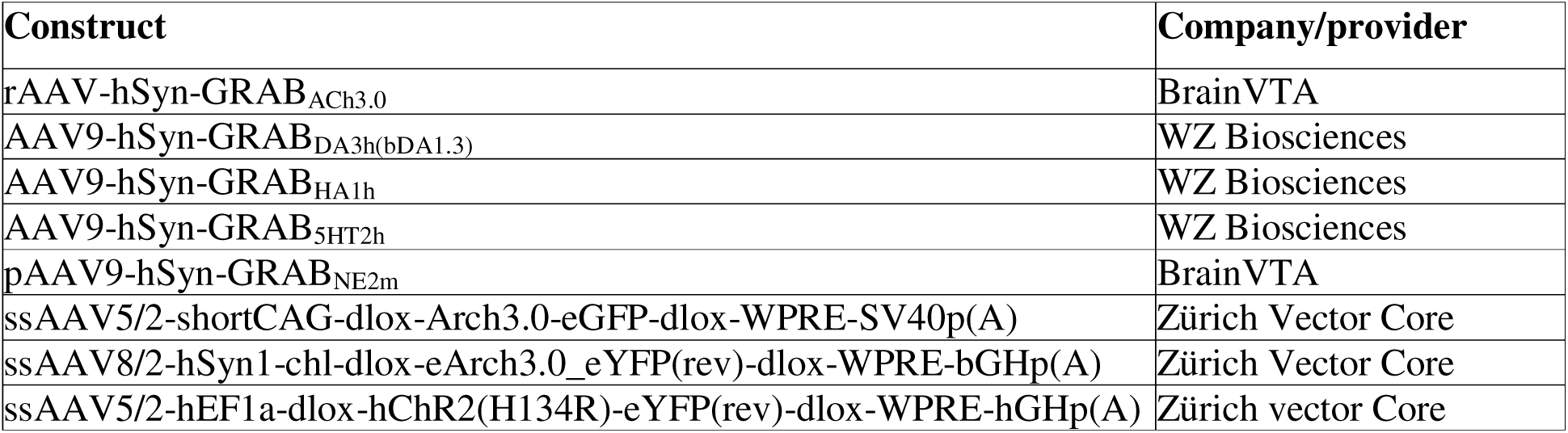
Viral constructs.

| Construct | Company/provider |
| --- | --- |
| rAAV-hSyn-GRAB<sub>ACh3.0</sub> | BrainVTA |
| AAV9-hSyn-GRAB<sub>DA3h(bDA1.3)</sub> | WZ Biosciences |
| AAV9-hSyn-GRAB<sub>HA1h</sub> | WZ Biosciences |
| AAV9-hSyn-GRAB<sub>5HT2h</sub> | WZ Biosciences |
| pAAV9-hSyn-GRAB<sub>NE2m</sub> | BrainVTA |
| ssAAV5/2-shortCAG-dlox-Arch3.0-eGFP-dlox-WPRE-SV40p(A) | Zürich Vector Core |
| ssAAV8/2-hSyn1-chl-dlox-eArch3.0_eYFP(rev)-dlox-WPRE-bGHp(A) | Zürich Vector Core |
| ssAAV5/2-hEF1a-dlox-hChR2(H134R)-eYFP(rev)-dlox-WPRE-hGHp(A) | Zürich vector Core |

**Table 2:** Primary antibodies.

| Name | Company/provider | Cat# |
| --- | --- | --- |
| GFP Polyclonal Antibody | Thermo Fisher Scientific | A-6455 |
| Anti-Tyrosine Hydroxylase, clone LNC1 | Merck Millipore | MAB318 |
| Anti-choline acetyltransferase (ChAT) | Abcam | AB178850 |
| Anti-GFP Monoclonal Antibody | Nacalai | GF090R |

**Table 3:** Secondary antibodies.

| Name | Company/provider | Cat# |
| --- | --- | --- |
| Goat anti-Rabbit IgG (H+L) Cross-Adsorbed Secondary Antibody, Alexa Fluor 488 | Thermo Fisher Scientific | A11034 |
| Goat anti-Mouse IgG1 Cross-Adsorbed Secondary Antibody, Alexa Fluor 568 | Thermo Fisher Scientific | A11004 |
| Goat anti-Rabbit IgG (H+L) Cross-Adsorbed Secondary Antibody, Alexa Fluor 568 | Thermo Fisher Scientific | A11011 |
| Goat anti-Rat IgG (H+L) Cross-Adsorbed Secondary Antibody, Alexa Fluor 488 | Thermo Fisher Scientific | A11006 |
| DAPI nucleic acid stain | Thermo Fisher Scientific | 62248 |

In order to conduct the optogenetic experiment targeting the locus coeruleus (LC) or nucleus basalis of Meynert, bilateral injections of AAV9/2 carrying floxed Arch3.0-eGFP, floxed hChR2-eYFP or control were administered. The injections were performed at the LC coordinates A/P −5.5 mm, M/L ±0.9 mm, and D/V −3.75 mm or nucleus basalis of Meynert coordinates A/P -0.35 mm, M/L ±1.6 mm, and D/V - 5.0 mm, -5.1 mm and -5.2 mm with a volume of 200 nl at each depth. Dual fiberoptic cannula (200 μm, 0.22 NA, Doric Lenses) was implanted above the LC at a depth of D/V −3.65 mm for LC targeting and mono fiberoptic cannulas (200 μm, 0.22 NA, RWD Life Science) were implanted bilaterally above the nucleus basalis of Meynert at a depth of D/V -5.1 mm for basal forebrain targeting.

### Fiber photometry

Each minicube (Doric Lenses) was connected to two pairs of excitation LEDs (465 nm and 405 nm, Doric Lenses, Tucker-Davis Technologies) using attenuator patch cords (400-μm core, NA 0.48, Doric Lenses). The minicube optics facilitated fluorophore monitoring through the use of dichroic mirrors and cleanup filters selected to match the excitation and emission spectra. LED drivers (Tucker-Davis Technologies, Doric Lenses) controlled the LEDs, which were linked to either a RZ10-X or RZ5 real-time processor (Tucker-Davis Technologies). In barrel cortex, bilateral excitation of biosensors was achieved by delivering 465-nm/405-nm light through separate patch cords on each side of the brain. The excitation at 465 nm/405 nm was sinusoidally modulated at frequencies of 531 Hz/211 Hz. Fiberoptic patch cords (400-μm core, NA 0.48, Doric Lenses or RWD Life Science) established the light pathway between the minicubes and the mice. Zirconia sleeves were utilized to attach the fiberoptic patch cords to the fiber implants on the mice.

A RZ10-X or RZ5 real-time processor was utilized to independently recover each of the four modulated signals generated by the four LEDs with a sampling rate of 1013. Standard synchronous demodulation techniques were employed for signal recovery. Synapse software (Tucker-Davis Technologies) facilitated control of the signal processor and alignment of fluorescent signals with video and EEG/EMG signals via incoming or outgoing TTL pulses.

MATLAB R2020a (MathWorks) was employed for data analysis. ΔF/F calculations of fluorescent signals were based on the fitted 405-nm signal or by using the median of the fluorescence signal itself. To obtain the scaled 405-nm channel, the 465-nm signal and the isosbestic 405 channel were first normalized using the least squares method (MATLAB polyfit function) to determine the necessary slope (a) and intercept (b):

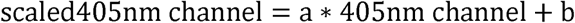

Subsequently, ΔF/F was calculated by subtracting the fitted control channel from the signal channel:

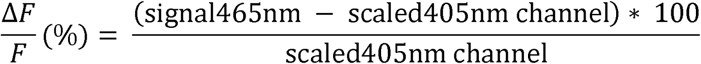

The power of infraslow neuromodulator fluctuations was done by extracting neuromodulator recordings from NREM including MAs lasting at least 120 s. The signal from each period was detrended and Welch’s method was used to estimate power. In the end, the weighed mean was calculated based on the period duration. The same periods were chosen for coherence analysis (*mscohere* function in MatLab), and the signals were downsampled by a factor 50 before analysis.

Cross-correlation analysis between continuous time series of norepinephrine levels and other neuromodulators was conducted during the following periods (lasting >120 s): NREM sleep including MAs, wake episodes>15 s and REM sleep. The calculation was done using the *xcorr* function in MatLab and the weighed mean was calculated based on the period duration.

State transitions were time-locked to norepinephrine troughs (the minimum value) during NREM sleep using custom-made Matlab code and divided into NREM-NREM, MA^EEG/EMG^ and wake^EEG/EMG^ depending on whether a sleep-scored MA and wake episode (see ‘Sleep measurement’) was present within 20 s of the norepinephrine troughs. ΔF/F values across transitions were normalized to span 0 to 1 for readily comparison independent of biosensor properties.

Frequency distributions of ΔF/F (%) of the neuromodulators were calculated for NREM sleep including MAs, wake episodes>15 s and REM sleep for the entire sleep session (3-4 hours) for all animals and divided into 0.5 ΔF/F bins.

### Sleep measurement

Mice were placed in recording chambers (ViewPoint Behavior Technology) with EEG and EMG electrodes connected via cables. These cables were then connected to a commutator (Plastics One, Bilaney, SL12C). The mice were given at least 24h to habituate to the recording chamber (ViewPoint Behavior Technology) before the actual recordings took place. On the recording day, the mice were attached to fiber-optic tethers, and recordings were conducted for 3-4 hours during their light phase. EEG and EMG signals were amplified (National Instruments, 16-channel AC amplifier, model 3500), and specific filtering settings were applied (EEG signal: high-pass at 1 Hz and low-pass at 100 Hz; EMG signal: high-pass at 10 Hz and low-pass at 100 Hz). Additionally, a notch filter at 50 Hz was used to minimize power line noise. Signals were digitized using a Multifunction I/O DAQ device (National Instruments, USB-6343) at a sampling rate of 512 Hz. Continuous video recordings were captured using an infrared camera (FLIR Systems) and later utilized for vigilance state scoring. Hypnograms were created by visually inspecting EEG traces divided into 5-second and subsequently 1-second epochs. Vigilance states were defined based on specific criteria related to muscle tonus and EEG characteristics; wake (high muscle tonus and a high-frequency, low-amplitude EEG), NREM sleep (no muscle tonus and low-frequency, high-amplitude EEG) and REM sleep (no muscle tonus and high-frequency, low-amplitude EEG). Wake bouts lasting less than 15 seconds were categorized as micro-arousals. Hypnogram analysis was performed using SleepScore software (ViewPoint Behavior Technology), while further data analysis was conducted in MATLAB using custom-made scripts. All data analysis was performed in MATLAB using custom-built scripts. Power spectral densities in the frequency range of 1–100 Hz were calculated using Welch’s method with a 5-second Hamming window, and the logarithm of the power was computed for each frequency. Average power was then calculated for specific frequency bands: delta (1–4 Hz), theta (4–8 Hz), sigma (8–15 Hz), beta (15–30 Hz), low gamma (30-60 Hz) and high gamma (60-80 Hz).

### Optogenetic modulation of LC and nucleus basalis of Meynert

A 532-nm laser (Changchun New Industries Optoelectronics Technology Co. Ltd., model 16030476) was utilized to deliver 2-minute continuous light exposure (5 mW at each fiber tip) bilaterally in LC or nucleus basalis of Meynert in order to activate Arch3.0, simultaneously with bilateral fiber photometry recordings in barrel cortex and EEG and EMG measurements. A three-hour closed-loop stimulation phase was initiated based on threshold crossing of smoothed real-time calculation of ΔF/F (%) calculations of NE levels based on 2 min windows (Synapse, Tucker Davis Technologies). This was done in order to achieve NREM sleep-specific stimulations and to compare similar brain states across animals rather than having random stimulations. A 465 nm laser (Shanghai Laser & Optics Century Co., Ltd., model: BL473-200FC) was used to generate 2 s stimulations of 10 ms pulses at 20 Hz (5 mW at each fiber tip) bilaterally in LC elicited by - 10 ΔF/F (%) threshold crossing of barrel cortex NE levels.

### Immunohistochemistry

To validate the location of the optic implant and confirm virus expression, we performed immunostaining on brain sections (Suppl. Figure S5A,D). Animals were deeply anesthetized with ketamine/xylazine and perfused transcardially with PBS, followed by 4% PFA. Brains were dissected, post-fixed in 4% PFA overnight, and transferred to PBS until sectioning. Using a vibratome, we cut 50-60 μm sections surrounding the implant sites. Sections were blocked at room temperature for 1–2 hours in PBS containing 5% goat or 10% donkey serum and 0.3% Triton X-100 before overnight incubation with primary antibodies at 4°C. After washing, sections were incubated with secondary antibodies for 2 hours at room temperature and brain slices were mounted on glass slides with DAPI-counting mounting medium.

Whole-brain slice images were acquired using Nikon Instruments Ni-E motorized microscope equipped with a 4× CFI Plan Apo Lambda objective (0.2 NA). Excitation was provided by a halogen light source with excitation filters for 362–389 nm, 465–495 nm, 530–575 nm, and Cy5 628–640 nm. Per section, 4×4 tiled images were captured and stitched automatically using Nikon NIS Elements AR software. For the nucleus basalis of Meynert, higher-magnification images were acquired using the same Nikon Instruments Ni-E motorized microscope with 10× and 20× CFI Plan Apo Lambda objectives (0.45 NA and 0.75 NA, respectively). The excitation sources, filters, and image acquisition and stitching settings remained unchanged for these higher-magnification images. For LC, higher-magnification images were obtained with a Nikon Instruments C2+ Ti-E confocal laser-scanning microscope, utilizing a 20× CFI Plan Fluor MI objective (0.75 NA) or a 40× CFI Plan Fluor oil objective (1.30 NA). Excitation sources included laser diodes at 405 nm, 561 nm, and 640 nm, as well as a 488 nm solid-state diode laser. For the 10× and 20x images, we acquired 80 Z-stacks every 1 um and flattened them (middle 20 ones out of 80) in Fiji/ImageJ using maximum intensity projection.

## QUANTIFICATION AND STATISTICAL ANALYSIS

Sample sizes were not determined using statistical methods, but they align with previous publications. Manipulations were randomized and counterbalanced across mice. Normality of data was assessed using the Shapiro-Wilk test. 1-way ANOVA or 2-way repeated measures ANOVA with Šídák’s multiple comparison post hoc test was used to assess grouped comparisons. Pearson’s correlation coefficient was used to assess correlation. All statistical test results are reported in the Supplementary Excel file.

## KEY RESOURCES TABLE

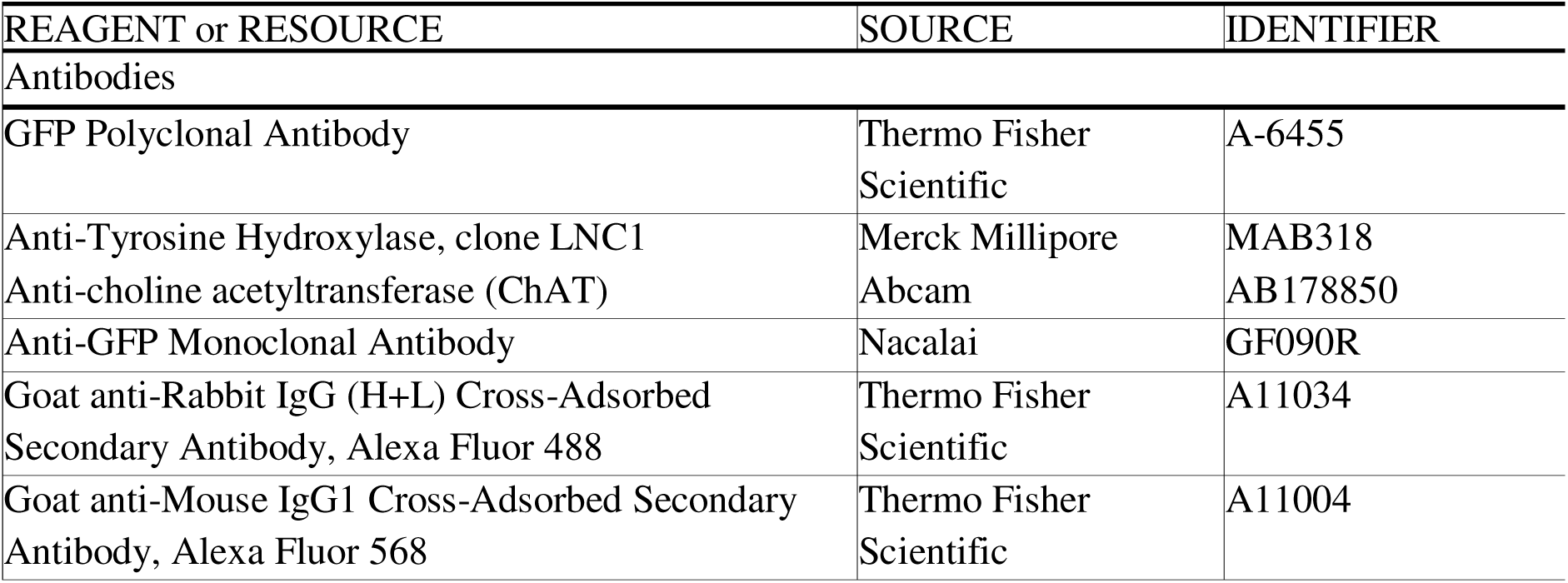
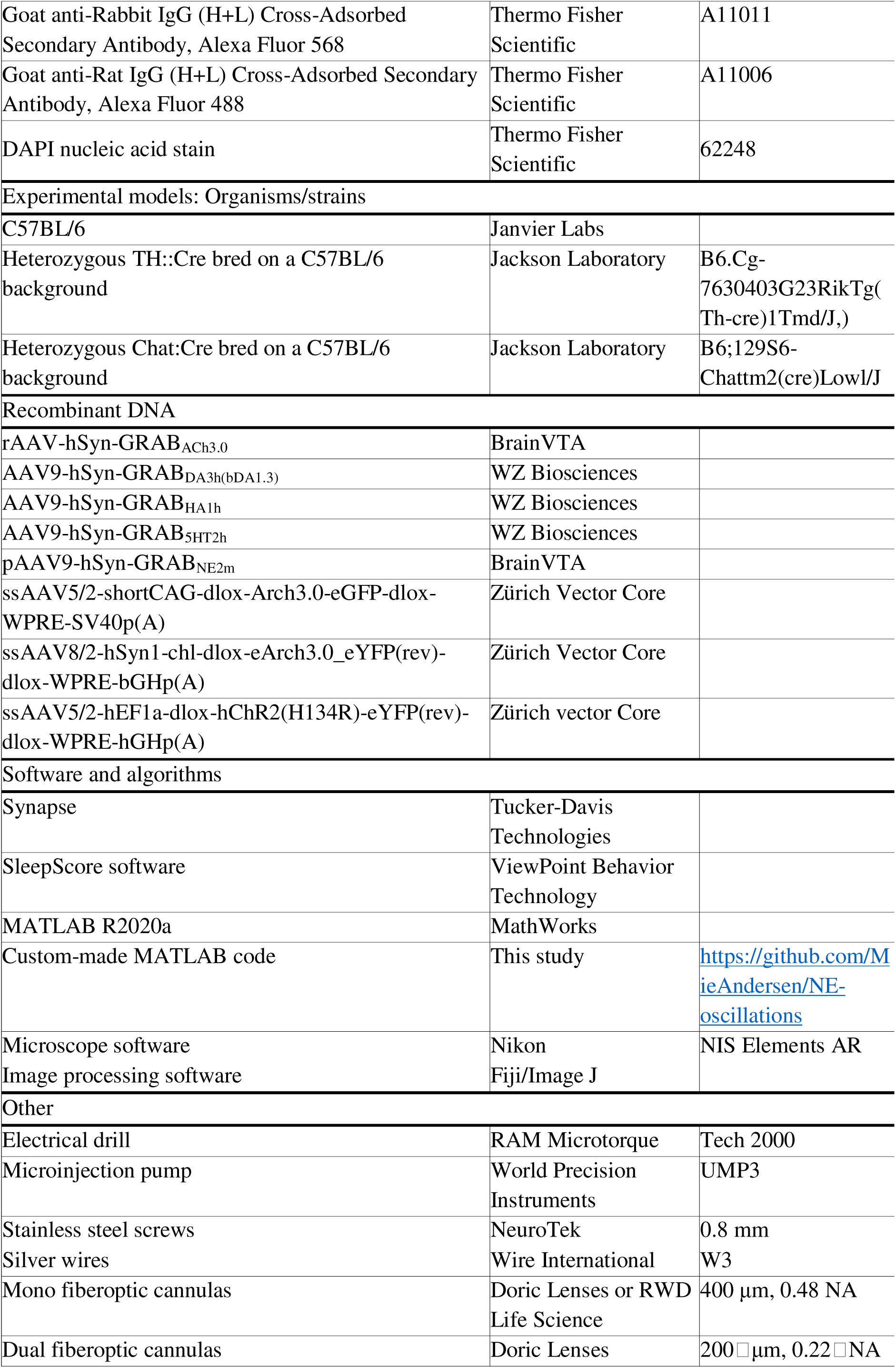
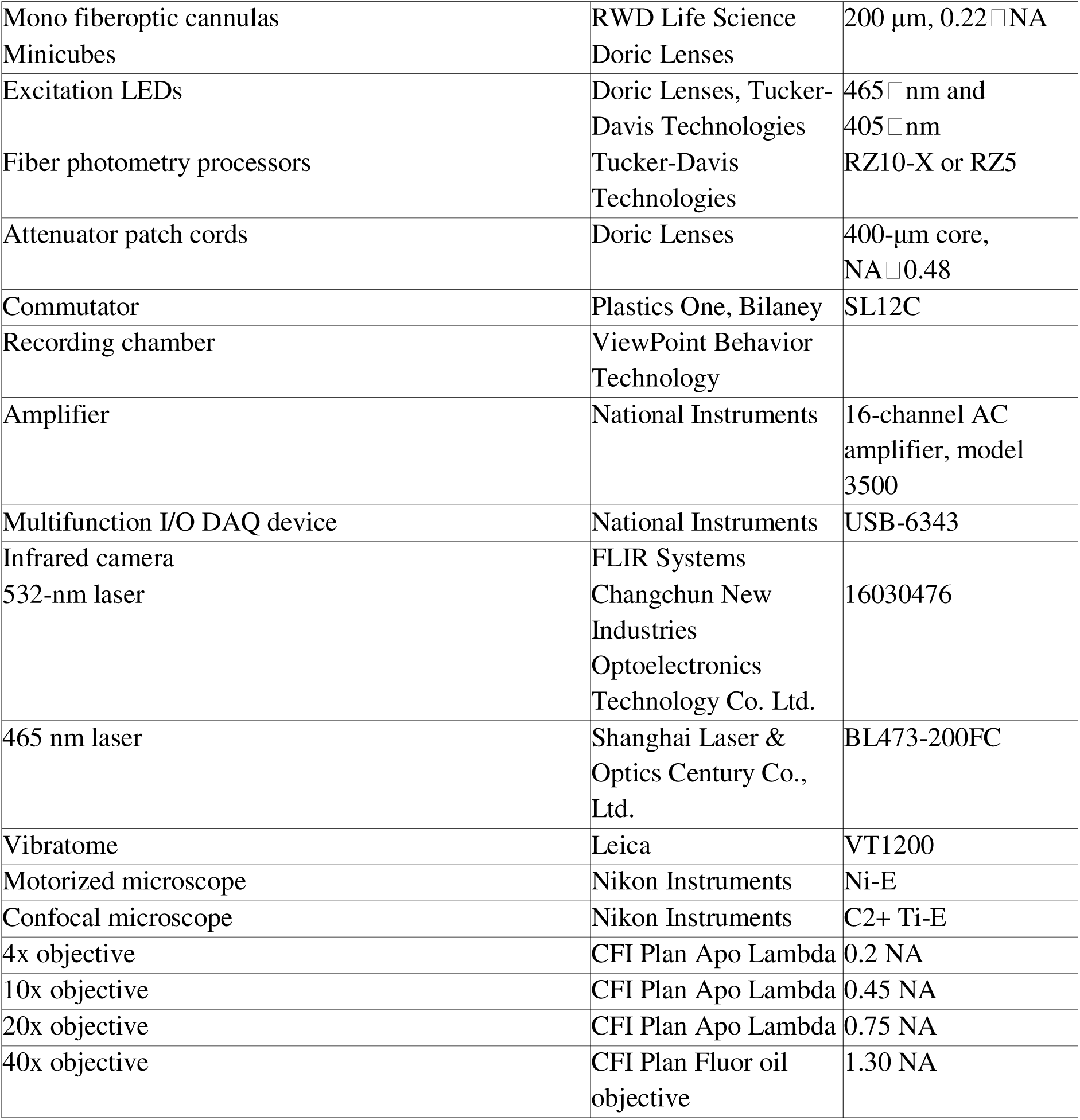

## SUPPLEMENTARY EXCEL FILE

**Table S1. Detailed results of statistical tests related to Figures 1-4, Figures S1-S3, and Figures S5-S7.** A.) Statistics related to Figure 1. B.) Statistics related to Figure 2. C.) Statistics related to Figure 3. D.) Statistics related to Figure 4. E.) Statistics related to Figure S1. F.) Statistics related to Figure S2. G.) Statistics related to Figure S3. H.) Statistics related to Figure S5. I.) Statistics related to Figure S6. J.) Statistics related to Figure S7.

## Notes

### Competing Interest Statement

The authors have declared no competing interest.

### Summary of Updates

This version of the manuscript has been revised to update missing figure numbers for the figure images in the uploaded pdf

