## Supplementary Figure for "Coordinated infraslow cortical oscillations of neuromodulators during NREM sleep"

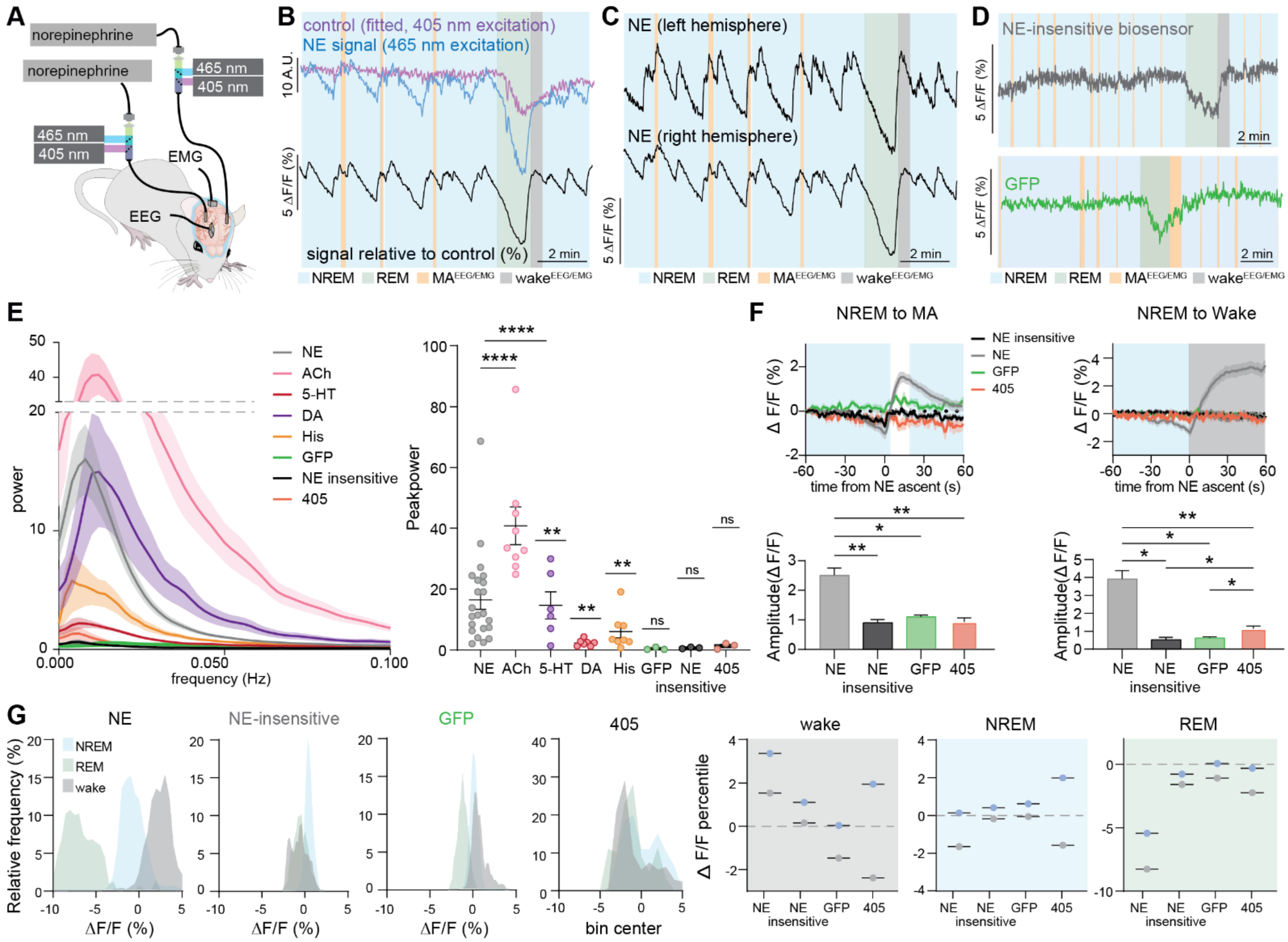

**Supplementary Figure S1. Control signals. Related to Figure 1.** (A) Each recording contained excitation of the target region with 405 nm light to get a control recording that contained norepinephrine (NE)-independent fluctuations. In addition, a group of mice was injected with the GRAB$_{NE2m}$ biosensor in both hemispheres of the primary somatosensory barrel cortex. (B) Example trace showing how the 405 nm control channel picks up artefacts during sleep. By using the 405 channel as the baseline for calculating the relative changes in fluorescence, these artefacts can be cancelled out (MA: micro-arousal). (C) Example traces of simultaneous NE (GRAB$_{NE2m}$) measurements from both hemispheres of barrel cortex illustrating the highly synchronous patterns of activity. (D) Example imaging traces of NE-insensitive GRAB$_{NEmut}$ (top) and green fluorescent protein (GFP) demonstrating tissue artefacts during sleep similar to appearance to 405. (E) Power of infraslow oscillations of neuromodulators, GRAB$_{NEmut}$, GFP, and 405 during NREM sleep and summary plots of peak power (Lognormal one sample t tests). (F) NE (GRAB$_{NE2m}$), NE-insensitive GRAB$_{NEmut}$, GFP, and 405 fluctuations shown across NREM-to-micro-arousal (MA$^{EEG/EMG}$) (top left) and NREM-to-wake (top right), as well as the corresponding amplitude of the increase (below corresponding plots) illustrating the higher response of the NE-sensitive biosensor (Mixed-effects model with Tukey's multiple comparisons test). (G) Frequency distributions as well as 25-75 percentile values of NE signal (GRAB$_{NE2m}$), NE-insensitive biosensor (GRAB$_{NEmut}$), GFP and 405 illustrating the much wider distribution of NE signal values across sleep and wake cycles compared to NE-insensitive and GFP controls (405 nm display wider distribution due to continuous photobleaching).

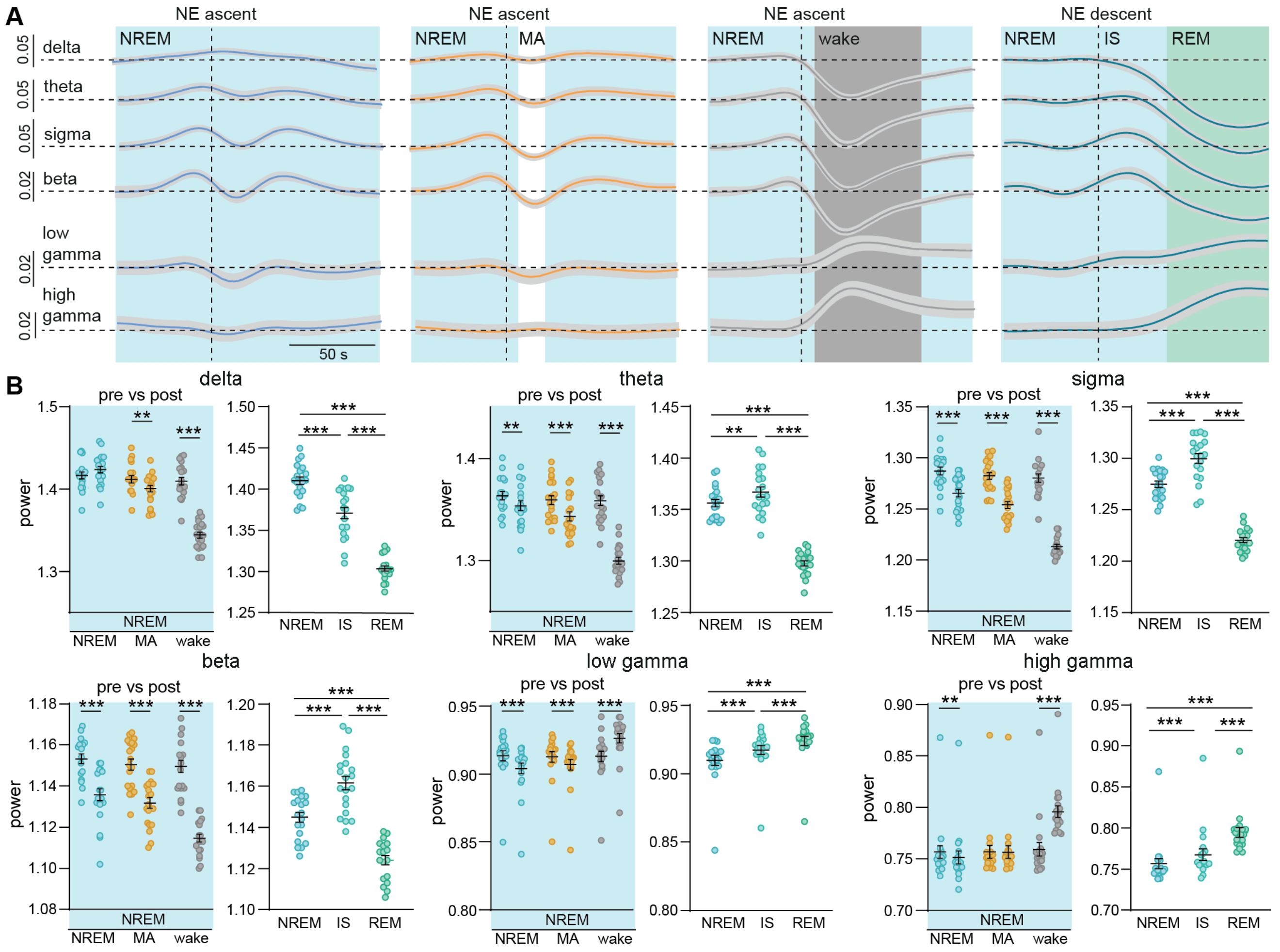


**Supplementary Figure S2. Spectral properties of sleep state transitions. Related to Figure 2.** (A) During NREM sleep, initiation of norepinephrine (NE) ascents was selected and divided into the following outcomes: 1) continuous NREM (blue), 2) micro-arousal (MA, orange) or 3) wake (grey). Furthermore, the initiation of NE descent preceding a REM sleep episode was selected and the period until REM sleep (green) onset was designated intermediate state (IS) sleep. The power of delta, theta, sigma, beta and low and high gamma were plotted prior to or following these time stamps (RM two- and one-way ANOVA or mixed-effects model). (B) Summary plots of the power of the different frequency bands pre- and post-transition. n = 20. Data is shown as mean±SEM. *$P$< 0.05, **$P$< 0.01, ***$P$< 0.001.

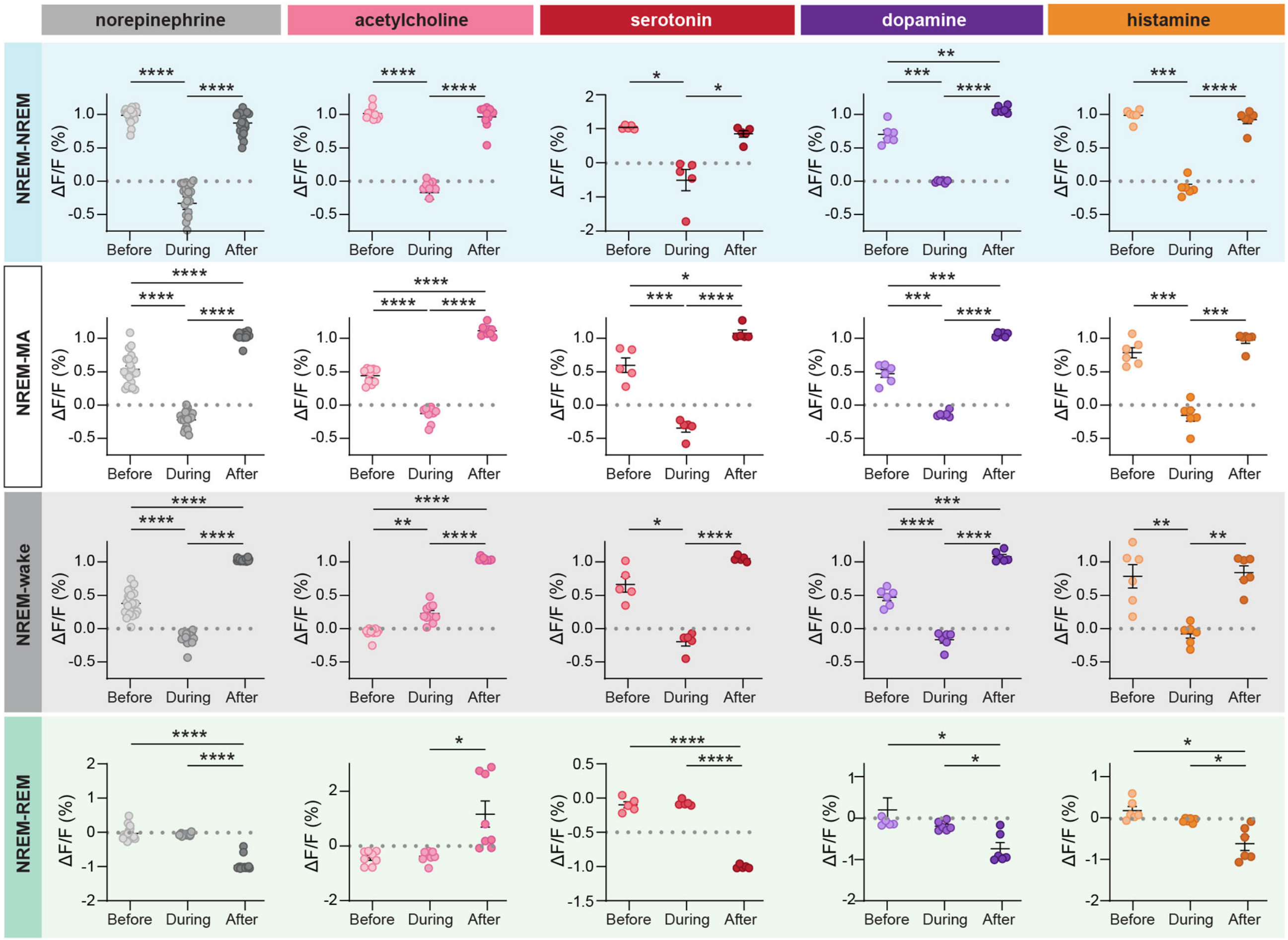


**Supplementary Figure S3. Summary plots of normalized neuromodulator fluctuation across transitions. Related to Figure 3.** Time stamps based on norepinephrine ascents resulting in continuous NREM sleep (blue), 2) micro-arousals (MA, white) or 3) wake (grey) and norepinephrine descents resulting in REM sleep (green) were used to extract the signal from the other neuromodulators and the signal normalized (max response = 1). Displayed are the summary plots of the extracted and normalized traces (from Figure 2b) (RM one-way ANOVA). Norepinephrine: n = 20; acetylcholine: n = 10; serotonin: n = 5; dopamine: n = 6; histamine: n = 6. Data is shown as mean±SEM. *$P$< 0.05, **$P$< 0.01, ***$P$< 0.001, ****$P$< 0.0001.

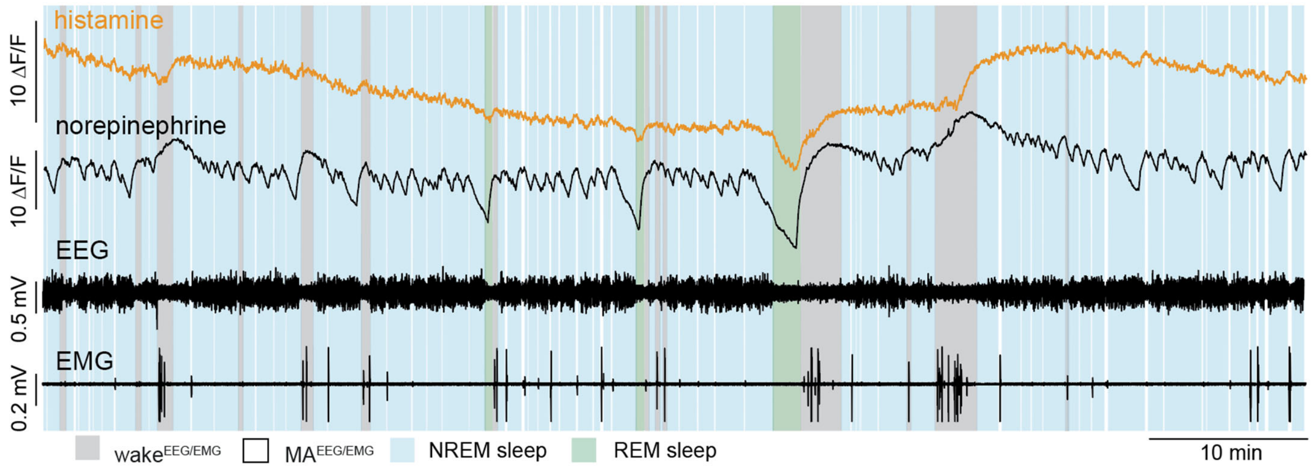


**Supplementary Figure S4. Slow dynamics of histamine. Related to Figure 4.** Example trace showing that in addition to the infraslow oscillations of histamine during NREM sleep, histamine also displays a gradual decline across NREM sleep independently of short microarousals (MA) and short awakenings. Upon longer awakenings, a large increase is observed, whereafter the gradual decline again happens.

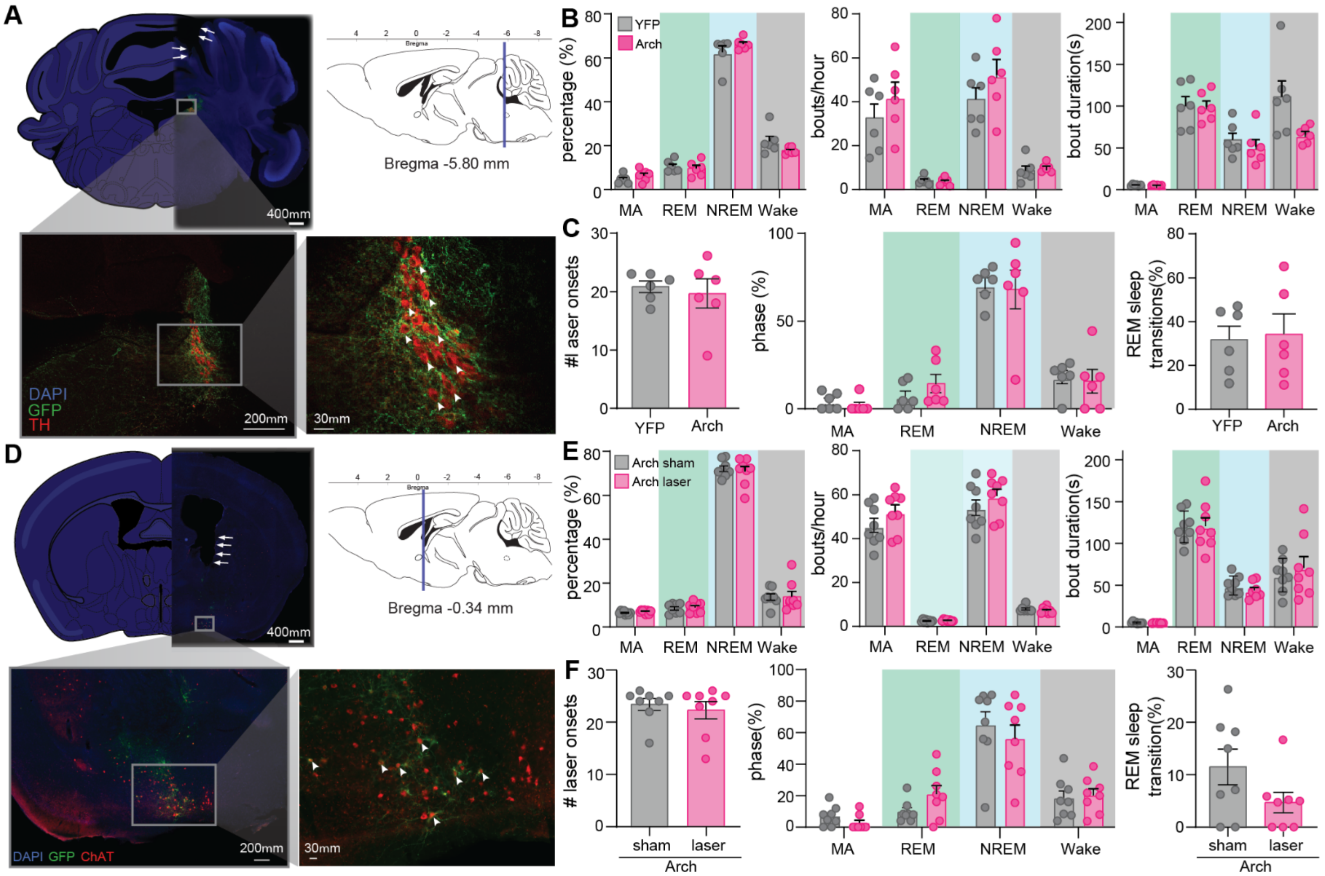


**Supplementary Figure S5. Optogenetic suppression of locus coeruleus and basal nucleus of Meynert. Related to Figure 3 and 4.** (A) Cre-dependent Arch (GFP) was expressed in tyrosine hydroxylase (TH) positive neurons in the locus coeruleus. (B) The overall composition of wakefulness, REM and NREM sleep as well as EMG-defined micro-arousals (MA) in animals expressing Arch and YFP in the locus coeruleus animals during laser on. (C) The mean number of laser onsets during each recording session as well as the brain state the laser onsets happen during. Last, the % laser onsets during NREM sleep that resulted in REM sleep transitions are shown (2-way repeated measures ANOVA with Šídák's multiple comparison post hoc test, unpaired t-test). (D) Arch (GFP) was expressed in choline acetyltransferase (ChAT) positive neurons in the basal nucleus of Meynert. White arrowheads show co-localized cells. White arrows indicate fiber placement. (E) Sleep composition in animals expressing Arch in basal nucleus of Meynert. Percentage of time spent in different stages, number of bouts per hour, bout durations (multiple paired t-tests with Holm-Šídák correction). (F) For optogenetic modulation of basal nucleus of Meynert is shown the number of laser onsets per sleep recording, the sleep state laser onsets happened in, as well as REM sleep inductions (paired t-test or multiple paired t-tests with Holm-Šídák correction). Locus coeruleus: n = 6 Arch, 6 YFP. Meynert: n = 8 Arch. Data is shown as mean±SEM.

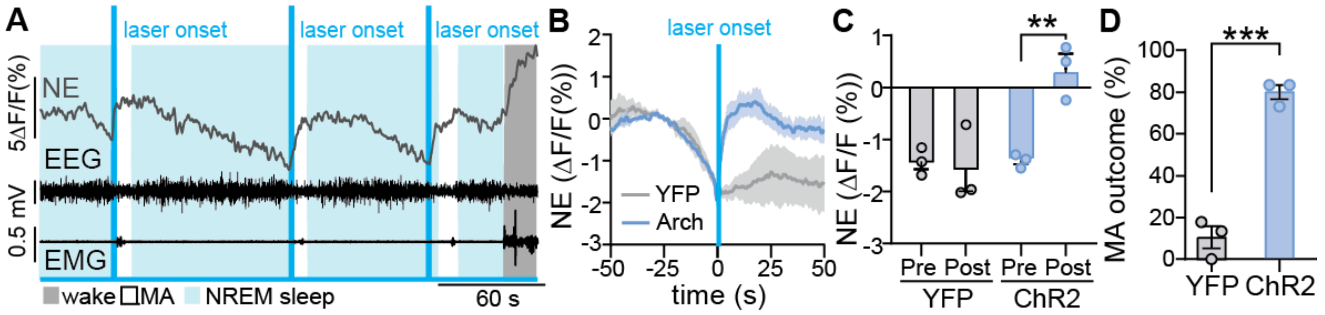


**Supplementary Figure S6. Optogenetic activation of locus coeruleus (LC) induces micro-arousals. Related to Figure 3 and 4.** (A) Example trace showing how optogenetic activation of noradrenergic neurons within locus coeruleus was initiated at norepinephrine (NE) declines > -10 ΔF/F (%) and led to immediate increases in NE in the barrel cortex. (B) Mean traces surrounding optogenetic activation of LC during NE declines during NREM sleep. (C) Summary plots showing NE prior to and after LC activation (Repeated measures two-way ANOVA with Šídák's multiple comparisons test). (D) The reported occurrence (%) of micro-arousals (MA) in the period following LC activation (0-20 s) in YFP and ChR2 expressing animals. YFP: n = 3; ChR2: n = 3. Data is shown as mean±SEM. $^{**}P < 0.01$, $^{***}P < 0.001$.

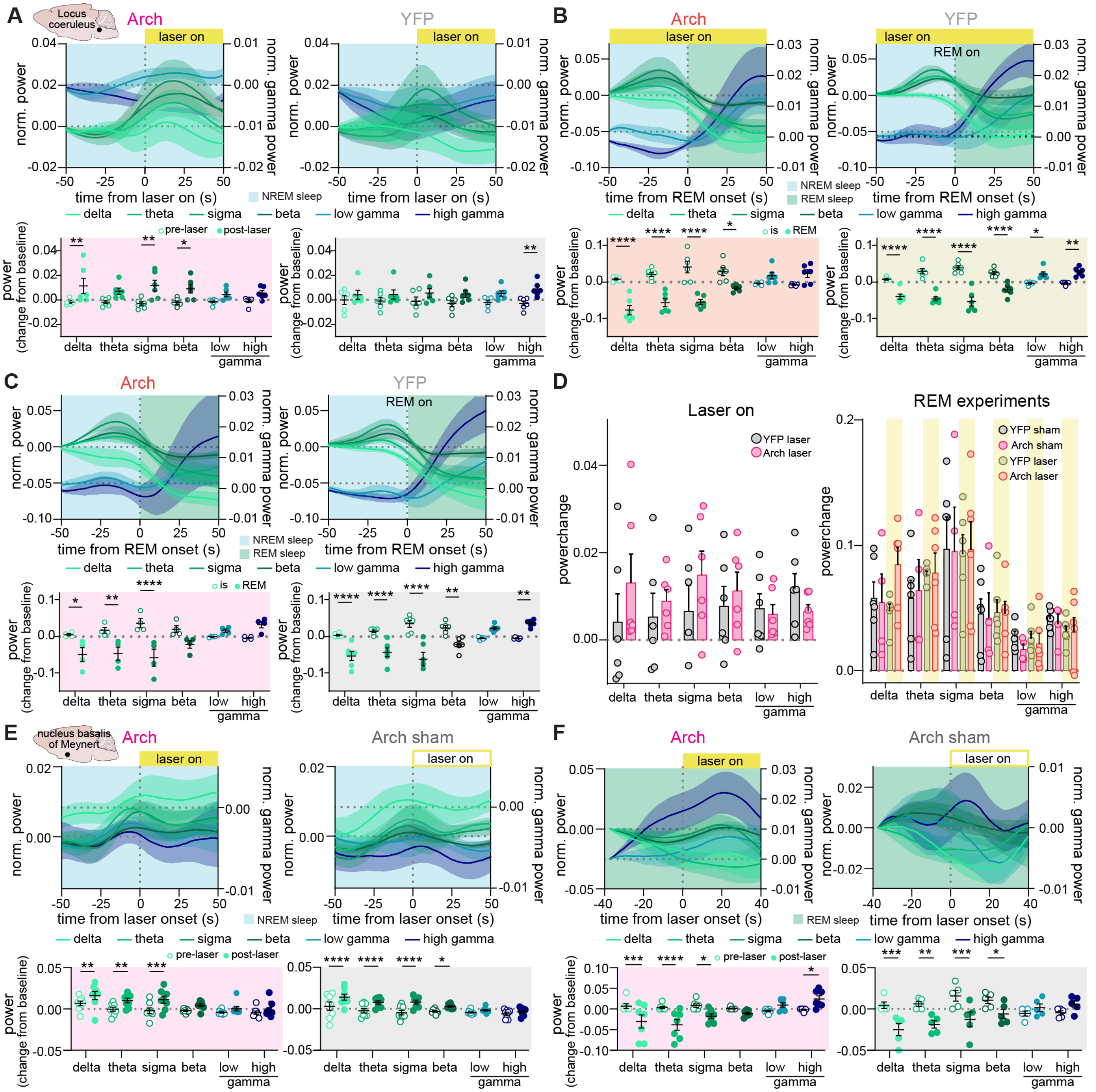


**Supplementary Figure S7. Spectral properties of EEG during optogenetic suppression of locus coeruleus. Related to Figure 3 and 4.** (A) Delta, theta, sigma, beta and low as well as high-gamma power shown before and after laser onsets during NREM sleep in animals expressing Arch or YFP in the locus coeruleus. Corresponding summary plots are shown below. (B-C) Spectral properties across REM sleep transitions for Arch and YFP animals when laser was on (B) versus off (C). Corresponding summary plots are shown below. (D) Summary plots showing the relative change in power across the NREM-laser on transitions (corresponding to a) for YFP and Arch animals and across REM sleep transitions (corresponding to b and c). (E-F) Spectral plots for suppression of basal nucleus of Meynert for laser on and off during NREM (E) and REM (F). All tests were done using 2-way repeated measures ANOVA with Šídák's multiple comparison post hoc test. Locus coeruleus: n = 6 Arch (REM Arch no laser = 4), 6 YFP. REM: n =

6 Arch, 6 YFP. Meynert: n = 8 Arch (REM Arch = 7, Arch no laser = 5). Data is shown as mean±SEM. **P* < 0.05, ***P* < 0.01, ****P* < 0.001, *****P* < 0.0001.
